# Texture coding in higher order somatosensory cortices of primates

**DOI:** 10.1101/2022.08.19.504511

**Authors:** Katie H. Long, Charles M. Greenspon, Ashley van Driesche, Justin D. Lieber, Sliman J. Bensmaia

## Abstract

Our sense of touch confers to us the ability to perceive textural features over a broad range of spatial scales and material properties, giving rise to a complex sensory experience. To understand the neural basis of texture perception requires that the responses of somatosensory neurons be probed with stimuli that tile the space of spatial scales and material properties experienced during everyday interactions with objects. We have previously shown that neurons in early stages of somatosensory processing – the nerves and somatosensory cortex (S1) – are highly sensitive to texture and carry a representation of texture that is highly informative about the surface but also predicts the evoked sensory experience. In contrast, the texture signals in higher order areas – secondary somatosensory cortex (S2) and the parietal ventral area (PV) – have never been investigated with a rich and naturalistic textural set. To fill this gap, we recorded single-unit activity in S2/PV of macaques while they performed a texture discrimination task. We then characterized the neural responses to texture and compared these to their counterparts in somatosensory cortex (S1). We found that the representation of texture in S2/PV differs markedly from its counterpart in S1. In particular, S2/PV neurons carry a much sparser representation of texture identity and also information about task variables, including the animal’s eventual perceptual decision. S2/PV thus seems to carry a labile representation of texture that reflects task demands rather than faithfully encoding the stimulus.

## Introduction

Our sense of touch confers to us a sensitivity to textures that spans six orders of magnitude in spatial scale (Skedung et al., 2013) and gives rise to a sensory experience that can span a vast space of qualities, such as roughness, hardness, stickiness, and warmth (Bensmaia & Hollins, 2003; Hollins et al., 1993; Lieber & Bensmaia, 2022). Previous studies with “natural textures” – which span the range of spatial scales and material properties experienced during everyday interactions with objects – have revealed that texture information is carried by multiple populations of nerve fibers using two distinct neural codes. Coarse textural features – measured in millimeters – are encoded in the spatial pattern of activation of one population of nerve fibers, which reflects the pattern of skin deformation produced by the surface. (Goodman & Bensmaia, 2017; Phillips et al., 2013). To perceive fine textural features – spanning tens of nanometers to hundreds of microns – requires movement between skin and surface, which elicits texture-specific vibrations in the skin; these vibration in turn drive highly repeatable and precise temporal spiking patterns in another population of nerve fibers (Bensmaia & Hollins, 2003, 2005; Greenspon et al., 2020; Long et al., 2022; Manfredi et al., 2014; Weber et al., 2013). These disparate neural codes – spatial and temporal – are reflected in the responses of neurons in somatosensory cortex (S1, including Brodmann’s areas 3b, 1 and 2), which reflect computations on spatial patterns of input activity to extract information about coarse textures and computations on temporal patterns of activity to extract information about fine features (Lieber & Bensmaia, 2019).

The highest known stages of processing along the medial lemniscus-dorsal columns pathway are the secondary somatosensory cortex (S2) and parietal ventral area (PV). Neurons in these areas receive projections from S1, have large receptive fields, often spanning one or even both limbs, and exhibit complex sensory response properties. Sensory responses in S2/PV are also more strongly modulated by selective attention than are their S1 counterparts (Gomez-Ramirez et al., 2014; Hsiao et al., 1993; Meftah et al., 2002). S2/PV neurons carry information not only about sensory stimuli but also about cognitive variables (Chapman & Meftah, 2005; Jiang et al., 1997; Romo et al., 2002; Rossi-Pool et al., 2021). For example, S2/PV neurons signal a change in texture when animals perform a texture discrimination task (Jiang et al., 1997) or the difference in frequency between two vibrations when they perform a frequency discrimination task (Romo et al., 2002). In other words, S2/PV responses reflect the decision variables that are relevant to the respective tasks. While these prior anatomical and neurophysiological studies were critical in establishing the position of S2/PV at the top of the touch neuraxis, neuronal responses were probed with stimuli that only varied one or two dimensions (vibratory frequency, orientation and curvature), which precluded an analysis of how naturalistic touch is encoded in these areas.

The goal of this study was to characterize the representation of a high-dimensional stimulus space in S2/PV, namely natural textures. Because S2/PV responses are dependent on the behavioral state of the animal and, in particular, are strongly modulated by the animal’s attentional state, we recorded neuronal responses while animals performed a texture discrimination task. We then assessed the degree to which S2/PV neurons carry information about texture and compared S2/PV texture responses to their S1 counterparts (Lieber & Bensmaia, 2019). First, we found that individual S2/PV neurons respond much more sparsely to texture than do S1 neurons. Second, the population response to texture in S2/PV has a different relationship to perception than does its S1 counterpart; indeed, the roughness signal – strong in S1 – is weak in S2/PV. Third, unlike their S1 counterparts, S2/PV responses do not carry texture information in precise temporal patterning. Fourth, S2/PV neurons encode information about task variables (whether the two textures presented on a given trial were the same or different) and about the animals’ eventual same/different judgment. We conclude that, while the elaboration of texture signals is modest in the first three stages of cortical processing (areas 3b, 1, and 2), these signals undergo a major transformation in S2/PV.

## Results

Two Rhesus macaques performed a ‘same-different’ texture discrimination task. On each trial, a rotating drum stimulator scanned two textures across the fingertip of the animal at a controlled speed (80 mm/s) and force (25 g) while the animal fixated its gaze on a screen (**Figure 1A**). At the end of the second texture presentation, the animal reported whether the two textures were the same or different by making a saccade to one of two targets (**Figure 1B**). A total of 44 diverse textures (including fabrics, furs, papers, see Supplemental Table 1) were presented in 90 unique combinations (45 ‘same’ and 45 ‘different’ pairs) while we recorded the neuronal activity using multi-contact electrodes (V-probes, Plexon Inc., **Figure 1C**). Texture presentations were counterbalanced such that each texture was as likely to be presented in the first as in the second interval. While animals performed the task, we recorded the responses of 158 neurons (across 3 hemispheres of the two animals) with receptive fields that included the fingertips (**Figure 1D**).

**Figure 1.**
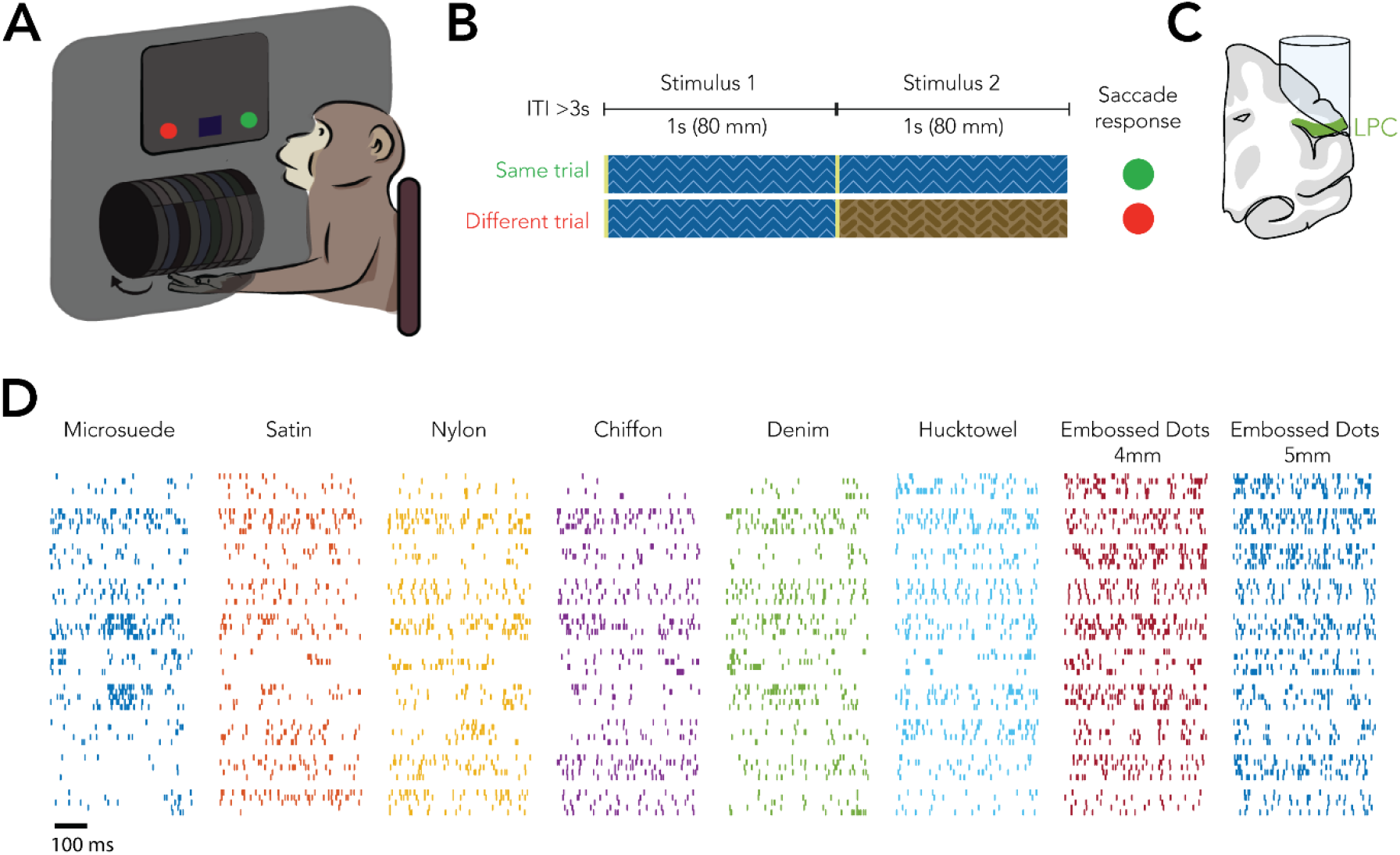
Experimental overview. A| The animal’s hand rested with the palm facing up on a hand-rest, configured to flex one finger so that the texture on the surface of the rotating drum only made contact with that fingertip. The animal’s view of the textures and its hand were obscured with a curtain. The trial progression was signaled via a computer monitor. B| On each trial, two textures were presented sequentially. The animal reported whether the two textures were the same or different by making a saccadic eye movement to one of two targets. C| Recording chambers were centered over the contralateral hand representation in lateral parietal cortex (over the upper bank of the lateral sulcus). D| Responses of 10 example S2/PV neurons (rows) to 5 repeated presentations of 8 example textures (columns). Different colors denote different textures.

### The discrimination of natural textures is a difficult task for monkeys

S2/PV responses have been shown to be strongly modulated by attention (Chapman & Meftah, 2005). With this in mind, we ensured that the animals were attending to the textured surfaces by having them perform a discrimination task. Despite extensive training over a period of around 2 years, the monkeys never achieved a high level of performance on this task. Indeed, performance leveled off at 69% and 55% across all texture pairs for the two animals, respectively (**Figure 2A**). The monkeys were more likely to correctly identify different pairs as different than they were same pairs (permutation test: p < 0.01 for both monkeys) and, accordingly, *d’*s were consistently positive across all texture pairs (**Figure 2B**). Monkey M (the better performing monkey) was relatively consistent across the 17 sessions (mean pairwise correlation of their behavioral choices = 0.93, p < 0.01) whereas Monkey S was considerably less consistent, though his behavioral choices were far more correlated across sessions than expected by chance (mean pairwise-correlation = 0.47, p < 0.01). Both monkeys thus attempted to perform the task: One monkey settled on a (suboptimal) strategy whereas the other adopted shifting and poorly performing strategies. Nevertheless, the monkeys’ attention was directed toward the textures.

**Figure 2.**
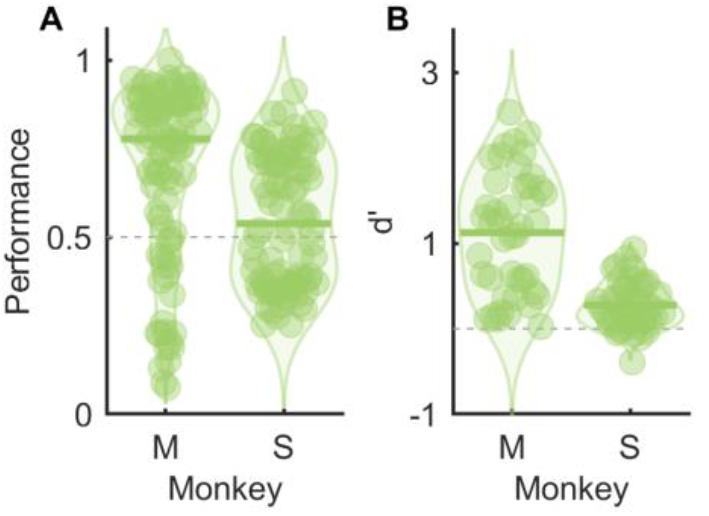
Behavioral performance. A| Proportion correct ‘same’ ‘different’ judgment for each pair of textures. B| Discriminability (*d’*) of the two textures in each different pair. Animals could discriminate different textures as evidenced by significantly positive *d’*s (mean *d’* = 1.12 & 0.28 respectively, permutation test: p < 0.01 for both animals).

Next, we investigated the basis of the animals same-different judgments. First, we assessed whether two textures were more likely to be judged as different to the extent that one was rougher than the other. We found that differences in perceived roughness – measured in human participants – were poor predictors of the tendency to judge two textures as different (**Supplementary Figure 1A, D**). Second, differences in population firing rate evoked by the two textures – a reliable neuronal correlate of differences in perceived roughness (Lieber & Bensmaia, 2019) – were poor predictors of the animals’ judgments (**Supplementary Figure 1B, E**). Finally, we assessed whether perceived dissimilarity or its neuronal correlate, neuronal distance (Long et al., 2022), could predict the same-different judgements and found that they could not (**Supplementary Figure 1C, F**). That is, we could not identify features of the textural percepts or of texture responses in S1 that could predict the animals’ judgments.

### The texture representation in S2/PV is sparse

Having established that the animals were engaged in the task, we gauged the texture sensitivity of S2/PV neurons and compared it to that of S1 neurons. First, we assessed how neurons responded to the range of textures. To this end, we gauged whether the response during the first interval significantly exceeded baseline. We focused on the first interval because the response in this interval was less liable to carry information about task variables (trial type, the animal’s decision). Of the 158 neurons, 120 (76%) responded significantly to texture, far fewer than in S1, where all 141 neurons did (**Error! Reference source not found.A, B**). Next, we assessed the sparseness of the texture responses, using as a gauge the proportion of textures that evoked a significant response in individual S2/PV neurons (using a criterion of 2 standard deviations above baseline). We found that, while most S1 neurons responded to most textures, even the most promiscuous S2/PV neurons responded to only a small fraction of all textures in the set (<20%, **Error! Reference source not found.C**).

This sparse representation of texture gave rise to a population-level representation of texture in S2/PV that was higher dimensional than its counterpart in S1. For the overlapping subset of textures, 21 principal components were required to explain 95% of the variance in texture responses in S2/PV whereas only 11 were required in S1. (**Figure 4A**; see **Supplementary Figure 2** for PCA over the full sets of textures). Similarly, response profiles were much more similar to one another for S1 than S2/PV neurons (r^2^ = 0.48 vs 0.18, **Figure 4B**).

**Figure 3.**
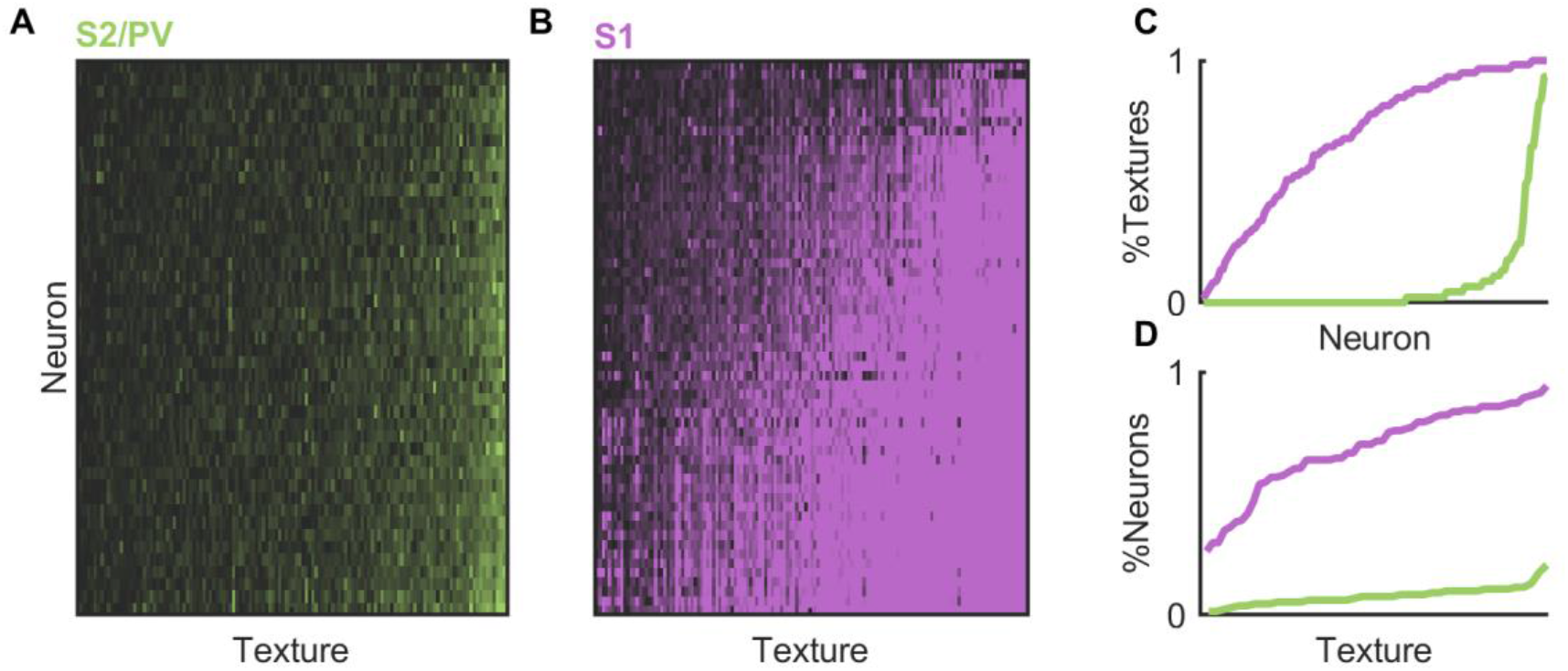
Texture responsiveness of S2/PV and S1. Z-scored responses of S2/PV (A) and S1 (B) neurons to various texture. Matrices are sorted both by number of textures that a neuron responds to (|Z| > 2, rows) and the number of neurons that each texture activates (columns). C| The proportion of textures that each neuron responds to (|Z| > 2) for both datasets. S2/PV neurons tend to respond to a lesser proportion of textures than their S1 counterparts. Similarly, (D) shows the proportion of neurons that each texture activates. We see that most textures only activate a minority of S2/PV neurons whilst in S1 most neurons are texture promiscuous. Note that both axes in (C) and (D) are normalized due to the different number of textures and neurons in each dataset.

**Figure 4.**
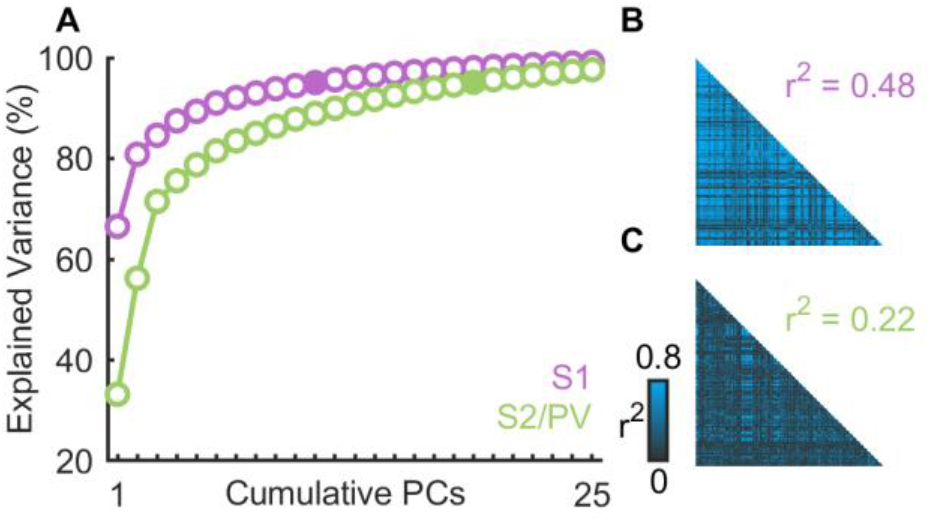
Dimensionality of the population level representation. A| Cumulative variance vs. number of PCs for S2/PV and S1 responses to matched textures. S2/PV is higher dimensional (n = 120, d = 21) than S1 (n = 141, d = 11) where d is the number of cumulative principal components necessary to reach 95% explained variance (filled circles). Correlation of the texture responses for pairs of neurons in S1 (B) and S2/PV (C). The responses of individual neurons to texture are more similar to each other in S1 than S2/PV. Results from the full texture set are shown in **Supplementary Figure 2**.

### S2/PV neurons carry weak information about texture

Next, we gauged the informativeness of S2/PV responses about texture. To this end, we first implemented a nearest neighbor classifier to assess the accuracy with which we could identify the texture presented on any given trial based on the responses of single units (**Figure 5A)**. We found that individual S2/PV neurons yielded poor classification performance (median performance = 3.3%, chance = 2.6%) compared to S1 neurons, which were individually more than twice as informative (7.3%, Wilcoxon rank sum: Z = -10.25, p < 0.01). Next, we examined the informativeness of the population response by performing a linear discriminant analysis with neuronal subpopulations of increasing size. Because the small number of trials obtained from each neuron impacted the performance of decoders applied to the much weaker and variable S2/PV response more than it did its S1 counterpart, we constructed a synthetic population for both S2/PV and S1 that allowed for a fairer comparison of decoder performance (see Methods and **Supplementary Figure 3**). We found that populations of S2/PV neurons were far less informative about texture than were their (size-matched) S1 counterparts (**Figure 5B**). While nearly perfect classification performance could be achieved with around 25 neurons in S1, the responses of 100 S2/PV neurons yielded a performance of around 40% correct, reflecting the sparseness, weakness, and variability of S2/PV responses (**Supplementary Figure 4**).

**Figure 5.**
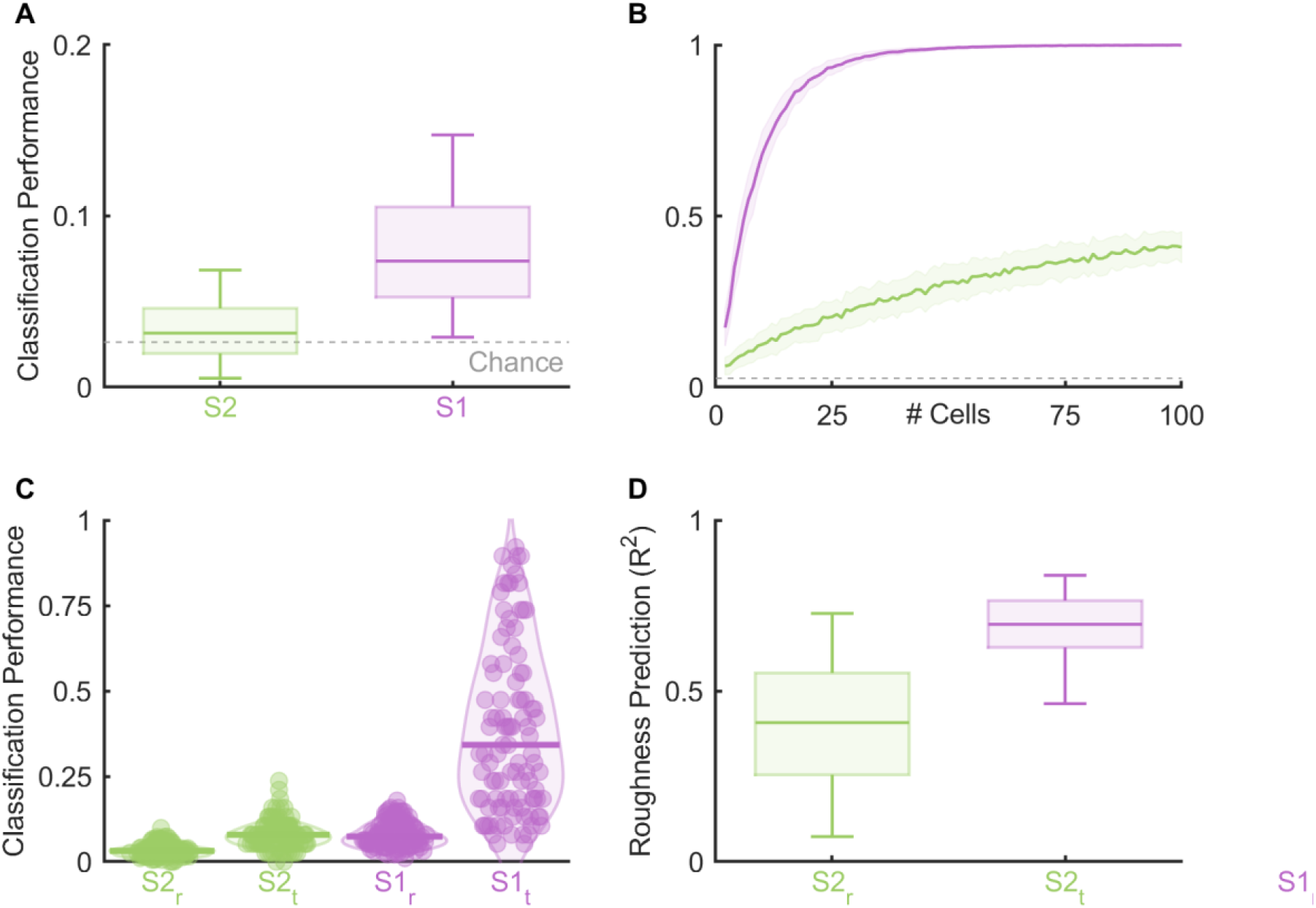
The texture representation is weaker in S2/PV than in S1. A| Performance of texture classifiers based on the responses of individual S2/PV and S1 neurons. Most S2/PV neurons (64%) and nearly all S1 neurons (95%) yield performance that is significantly above chance (64%). Individual S1 neurons are far more informative than are their S2/PV counterparts. B| Performance of texture classifiers based on the population response in S2/PV and S1 neurons as a function of the population size. Near perfect classification performance is achieved with around 25 S1 neurons. In contrast, performance peaks at 38% with the entire S2/PV sample. Lines represent the median while shaded area indicates 25^th^ and 75^th^ percentiles. C| Individual S2/PV neurons are poor predictors of perceived roughness (median R^2^ = 0.41) in contrast to their S1 counterparts (median R^2^ = 0.71). D| Timing information is present in S2 neurons and enables improved classification at a single cell level though the improvement is minor in comparison to S1. Subscript indicates if a rate (r) or timing (t) classifier was used.

### Temporal spiking patterns are largely uninformative in S2/PV

Tactile nerve fibers have been shown to carry texture information in their temporal spiking patterns. Indeed, scanning a textured surface across the fingertip (or vice versa) produces vibrations in the skin whose properties depend on the scanned texture (Bensmaia & Hollins, 2003, 2005; Greenspon et al., 2020; Manfredi et al., 2014). These vibrations in turn produce millisecond-precision temporal spiking patterns in nerve fibers that carry information about texture, particularly about fine textural features (Weber et al., 2013). This temporal code for texture is also observed in a subpopulation of S1 neurons (Lieber & Bensmaia, 2019; Long et al., 2022). With this in mind, we examined the degree to which textures could be identified based solely on the temporal patterning in the S2/PV response. To this end, we performed a classification analysis based on the temporal spiking patterns, excluding any rate based information. Classification performance was high to the extent that temporal spiking patterns evoked by repeated presentation of one texture were similar to each other and differed from the patterns evoked by other textures. We found that texture classification based on spike timing was better than chance but far inferior to that achieved with S1 neurons (**Figure 5C**). We conclude that, while some temporal patterning is preserved in S2/PV, this patterning is far less informative than is that in S1.

### S2/PV responses are poor predictors of roughness

Roughness is the dominant axis of texture, accounting for a large fraction of judgments of how dissimilar two textures feel (Bensmaia & Hollins, 2005; Hollins et al., 2000). S1 responses have been shown to be highly predictive of the roughness of a texture. Indeed, the first principal component in the S1 population response is a strong predictor of perceived roughness (r^2^ = 0.72). With this in mind, we examined whether this roughness signal was also found in S2/PV. To this end, we regressed the first principal component of the S2/PV population response on judgments of roughness of the corresponding textures, obtained from human observers. We found that the first PC in the neuronal response was only weakly correlated with roughness (r^2^ = 0.2) but the second one was more so (r^2^ = 0.45). We then assessed whether the roughness signal was present but distributed more widely in S2/PV than in S1 by regressing roughness on the responses of individual neurons (**Figure 5D**). Even then, roughness predictions based on S2/PV responses were far poorer than those based on S1 responses (R^2^ = 0.41 vs. 0.71 for S2/PV and SC, respectively). Thus, roughness, the dominant axis of texture perception, is less reliably and more diffusely represented in S2/PV than in S1.

### S2/PV responses carry information about task variables

S2/PV responses have been shown to carry not only sensory information but also information about task variables. For example, in a two-alternative forced choice frequency discrimination task, S2/PV responses to vibrations were proportional to the difference in the frequencies of the first and second vibrations, the quantity relevant to the animals’ ultimate judgment (Romo et al., 2002). In a texture change detection task, a subset of S2/PV neurons responded preferentially to texture changes, again carrying the outcome of the computation relevant for the animals’ ultimate judgment (Jiang et al., 1997). In light of this, we examined whether these task variables were also observed in the S2/PV responses when animals performed a ‘same-different’ task. To this end, we compared S2/PV responses on same vs. different trials, focusing on the second interval, when the trial identity becomes apparent. Given that textures were counterbalanced and equally likely to appear in the first or second interval regardless of trial type, divergence of the response in the second interval would indicate that neuronal responses to the same texture in the second interval would depend on whether or not that texture was also presented in the first interval. We found that the responses of some neurons indeed diverged in the second interval when conditioned on trial type (**Figure 6A**, top). When we conditioned the responses on the animals’ ultimate decision (same vs. different), we again found the responses of some neurons to diverge in the second interval (**Figure 6A**, bottom). For these neurons, the response to a texture on a ‘same’ trial resembled that on a ‘different’ trial if the animal reported these two textures as being the same. Most neurons showed a combination of these response properties. Across the S2/PV population, 54% encoded trial type only, 26% of the neurons encoded the animal’s decision only, and 15% encoded both trial type and the animals’ decision (t-test, p < 0.01). Note that the majority of these neurons also carried information about texture. To assess these task signals across the population, we used a linear discriminant analysis to classify trial type and decision as the trial progressed (**Figure 6B**). We found that we could predict the trial type soon after the onset of the second texture. This trial-type signal was phasic and decreased to near chance at the end of the second interval. The emergence of the trial-type information was followed shortly by the emergence of a decision signal which increased until the animals’ eventual decision.

**Figure 6.**
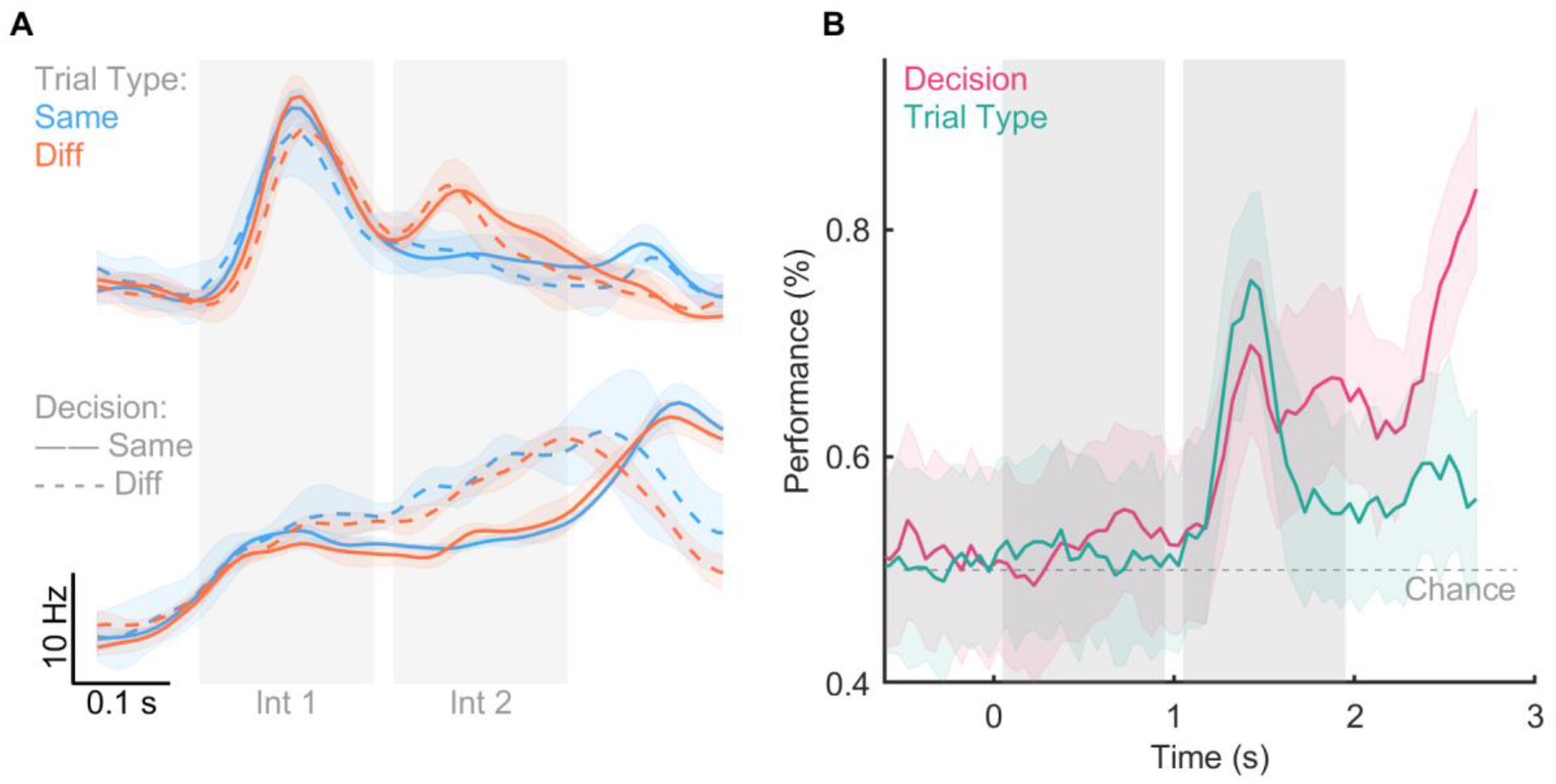
Task-related signals are present in S2/PV. |A| Peri-stimulus time histogram of two example neurons which differentially respond to trial type (top) and the animal’s decision (bottom). B| Sliding window classification of both the animal’s decision and the trial type. Line color indicates trial type and line style indicates the animal’s decision. Line indicates mean and shaded areas indicate standard error of the mean. Shaded panels indicate the stimulus intervals (B) Classification performance for sliding window LDA classifier of trial type or decision (chance = 0.5). Line indicates mean while shaded area indicates standard deviation.

## Discussion

### S2 neurons carry sparse texture signals

Neurons in S1 and S2/PV differ starkly in their overall responsiveness to texture. Whereas almost all neurons in S1 respond to texture and tend to respond to most textures, many neurons in S2/PV do not respond to texture at all, and those that do respond only respond to a small number of textures. The increased sparseness is consistent with observations in other sensory pathways, where dense coding of low-level features in early stages of processing gives way to sparse activation and increased selectivity in higher order cortical fields (Chalk et al., 2018; Okazawa et al., 2017). The sparseness of the texture representation in S2/PV is one of the factors that leads to a less informative representation of texture than in S1, as evidenced by poorer texture classification performance. To the extent that individual neurons only respond to a small fraction of textures, a relatively small sample of such neurons will fail to encode some textures. A second contributing factor is that the variability of S2/PV responses is higher than its counterpart in S1, as evidenced by a Fano factor that is almost twice as high. One source of this variability may be the multiplexing of information about texture and task variables, largely absent in S1 (see below).

### S2 neurons encode task variables

Across behavioral conditions, a subpopulation of S2/PV neurons has been shown to be modulated by task-relevant variables. In a texture change-detection task, the responses of most neurons indicated the presence, but not magnitude, of a change in texture (Jiang et al., 1997). In an active texture exploration task, many neurons were anecdotally found to be responsive to a small metal bar that demarcated the beginning, middle, and end of the texture (Sinclair & Burton, 1993). In a vibratory frequency discrimination task, a subset of neurons encoded the difference in frequency of two, sequentially presented vibratory stimuli, the decision variable relevant to the eventual decision (Romo et al., 2002). In a vibratory same-different task, a subset of neurons encoded the trial-type (same, different), as we found here (Rossi-Pool et al., 2021). We found that information about the key task variables – trial type and the monkey’s decision – were encoded in S2/PV. However, decision- and task-related signals were multiplexed with texture-related ones, such that information about all these task variables could be decoded from the population response. These different signals were not segregated across different populations of neurons, as evidenced by the fact that neurons were often identified as responding to more than one feature (texture, decision, change detection).

### S2 neurons only weakly encode roughness

S1 neurons carry a large, shared signal (roughness) – which accounts for two thirds of the variance in the population response and is highly predictive of perceived roughness. The population response in S2/PV is not dominated by a single dimension, and the axis of highest variance in S2/PV responses is a poor predictor of roughness. In fact, the entire S2/PV population response yields poor predictions of roughness. One possibility is that roughness signals were weak in S2 because the animals did not base their same-different judgments on roughness. Indeed, differences in perceived roughness (measured in humans) were poor predictors of the animals’ tendency to judge two textures as different. From this perspective, S2 would carry a robust roughness signal if the animals were performing a roughness discrimination task. Another possibility is that S2/PV neurons encode higher level texture attributes at the expense of low-level features like roughness. Consistent with this hypothesis, several neurons exhibited a preference for animal fur, a high-level attribute. The texture set was too small to explore selectivity for specific high-level textural features. Nevertheless, the progression from neurons that encode low-level stimulus features to neurons that encode higher-order ones as one ascends the neuraxis is observed in other sensory modalities (Groen et al., 2017; Rauschecker, 1998; Yamins & DiCarlo, 2016).

## Conclusion

S2/PV carries a sparse representation of texture in which signals about low-level stimulus features such as roughness have been strongly attenuated, perhaps replaced by signals about high-level features. The sensory signals are multiplexed with cognitive ones.

## Methods

### Animals

All experimental procedures involving animals were approved by the University of Chicago Institutional Animal Care and Use Committee. Behavioral and neurophysiological data were obtained from two rhesus macaques (males, 10-14 years old, 8-10 kg). Both animals were instrumented with a custom head-post to allow for head immobilization for eye-tracking and stable neurophysiological recordings. After two years of head-fixed training on a behavioral task (described below), animals were instrumented with a 22-mm diameter recording chamber (Crist Instrument Co., Hagerstown, MD) positioned over the upper bank of the lateral sulcus (Figure 3.1C). In both animals, chambers were centered 8-10 mm anterior to ear bar zero, as far lateral as possible, and positioned straight up and down using a steep chamber angle (30-45°) to keep the edge of the chamber flush with the skull. Coordinates were determined using a stereotaxic instrument (David Kopf Instruments, Tujunga, CA). In one hemisphere of one monkey, a flat 0° chamber was placed flush with the skull, such that it was positioned at a ∼40° angle centered over the lateral sulcus. The chamber was placed at this modified angle to accommodate the placement of a 96-channel Utah array (Blackrock Microsystems, Salt Lake City, UT) over the hand representation in Area 1.

### Neurophysiology

Each day, extracellular responses were obtained using one or two 16-channel linear arrays (V-probes, Plexon Inc, Dallas, TX). Electrodes were moved using a computer-controlled microdrive (NAN Instruments, Nazaret Illit, Isreal) and responses were recorded through a Cereplex Direct and Central software (Blackrock Microsystems, Salt Lake City, UT). Spike sorting was performed manually using Offline Sorter (Plexon Inc, Dallas, TX) in order to identify well-isolated single units that were stable across the recording session.

### Behavioral Task Task structure

Animals were trained to perform a 2-alternative forced-choice task in which they reported whether two sequentially presented textures were the same of different. Two texture stimuli were presented sequentially, each for 1s, with no interstimulus interval separating them. Same trials began at the start of a given texture strip, such that the full 16cm were scanned across the finger over a 2s period. A thin strip of tape (3 mm wide) placed at the beginning and halfway through each texture (8 cm) demarcated the start of each stimulus period. On different trials, the stimulus pair began at this halfway point on the first texture and moved halfway through the subsequent texture on the drum. The drum never lifted off the fingertip during the trials, and there was no gap between textures. This design was chosen because a delay period between the stimuli increased the task demands beyond our animals’ capabilities. However, it also constrained our stimulus set such that any given texture was, by necessity, always followed by one specific texture rather than a variety of possible comparisons. Following the 2 s trial, the animal made a saccade toward a right or left target to indicate whether the two textures were the same or different, respectively.

45 unique textures were presented in 90 pairs for 5 repeats across four experimental blocks. During the first block, 40/90 texture pairs were presented three times each in pseudo-random order. During the second block, the remaining 50/90 texture pairs were presented three times each in pseudo-random order. The third and fourth blocks followed the same design but contained only two repeats of each stimulus pair.

### Texture Stimuli

Textures were scanned across the fingertip at 80 mm/s and 25 g of force. Textures were presented using a custom stimulator that included a rotation motor (SmartMotor SM23165D; Animatics) connected to a 1:100 gearbox (Animatics), which provided precise control of rotational position (±200 μm) and velocity (±1.1 mm/s). A vertical stage (IMS100V; Newport) controlled the depth of indentation into the skin with a precision of 2 μm. This rotating drum was also connected to a horizontal stage (IMS400CCHA; Newport), allowing for horizontal displacements over a range of 40 cm (±4 μm). With this rig, we maintained precise horizontal, vertical, and rotational positioning of textures. We presented 44 unique textures, including fabrics, furs, papers, and 3 3D-printed stimuli. We also included a 45th stimulus as a catch trial in which the drum moved like in any other trial, but no texture contacted the fingertip. 40 of these 44 textures were also used in our previous examination of texture coding in anterior parietal cortex, and only responses to these 40 textures were used for all comparisons of S1 and S2/PV.

### Somatosensory Cortex Responses

All S1 data were collected from awake macaques that were performing a visual contrast-discrimination task. All stimuli were presented as described below. However, unlike in the present study, only one stimulus was presented in each trial, with at least 3 seconds between trials. Further details on these methods can be found in previous reports on these data (Lieber & Bensmaia, 2019; Long et al., 2022).

### Human Psychophysical Ratings

We leveraged a dataset on the perceived roughness of the 40 textures in our shared SC-S2/PV stimulus set. We have already published reports using these data (Delhaye et al., 2019; Lieber & Bensmaia, 2019). Briefly, we asked six human subjects (5 males, 1 female, ages 18-24) to freely rate the perceived roughness of textures presented in the same manner as described here (80 mm/s, 25 g). All textures were presented six times to each subject and ratings were normalized across experimental blocks and across subjects.

### Data Analysis

All analyses and statistics were performed using Matlab 2022a (Mathworks, Natick, MA, USA).

### Texture pair discriminability

The discriminability value for a given AB different pair was computed by first determining the false alarm rate for each texture separately – the rate at which the AA or BB stimulus pairs were reported as different – taking the average of the false alarm rates, and then comparing the hit rate – the proportion of trials in which the AB stimulus pair was correctly identified as different. This was computed across all days after behavioral performance had stabilized. Values were then converted to their Z-score probabilities (normal inverse function) and then the false alarm rate was subtracted from the hit rate.

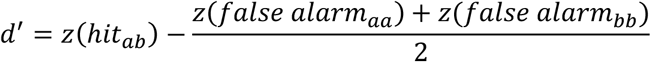

### Behavioral strategy

For each animal we took the proportion of trials for which a given texture pair was selected as different. We then randomly split the sessions in half and combined the proportions for each half. Then, we computed the correlation between half of the sessions to determine how similar one randomly selected half was to another. We repeated this bootstrap 1000 times to get a mean similarity rating. To determine significance, we repeated the above procedure but shuffled the pair identities before computing a null correlation. The observed mean value was then compared to the null distribution and the proportion of bootstraps for which the observed value was smaller reported as the p-value.

### Firing rates & peri-stimulus time histograms

Baseline firing rates were calculated as the mean firing rate (spikes/s) over a 600 ms window prior to the texture contacting the skin. Texture-evoked firing rates were calculated over a 300 ms window following the texture-evoked transient during each stimulus interval. The decision window was 600 ms starting 200 ms after the end of the second stimulus. Texture evoked firing rates were used for all further analyses unless stated otherwise.

For PSTH production, spike rates were computed for 5 ms bins. Error bounds were computed by randomly allotting the trials into 5 folds and then computing the standard deviation of the folds at each time point. Finally, a 25 ms gaussian smoothing filter (5 bins wide) was applied to the PSTHs, though smoothed data was never used for analysis.

### Cell responsiveness

Two criteria were used to determine if a given neuron was responsive to texture or not and if either criterion was met then the neuron was included. First, across all trials the baseline and first interval firing rates (post-transient) were compared with a paired t-test. Due to the sample size and number of cells the alpha was set to 1e^-4^ to minimize false positives (note that, for the number of neurons, the Bonferroni-adjusted alpha would be greater than 1e^-4^). Separately, we also performed a permutation test to determine if the neuron was responsive to a subset of textures that would prevent detection with the paired t-test. For each texture we took the mean evoked response to a texture when it was presented in the first interval (across same and different pair trials). We then built a null distribution (n = 10,000) of differences between randomly selected baseline and interval 1 firing rates and compared the mean evoked response to the null distribution. If any of the textures were considered significant (p < 0.05) we included the texture. Roughly 53% of neurons met both criteria, 20% just met the first, 10% just the second, and 17% neither.

Similar to the above, to determine if each cell encoded the animal’s decision or acted as a change detector, we split the responses during the second interval or post-stimulus interval into same/different trials or chose same/chose different trials respectively. We then performed a 2-sample t-test for on the appropriate subsets of trials and identified a neuron as being sensitive to one or both depending on the results of the test with an alpha of 0.005.

### Skew

To determine whether or not the neurons selectively responded to given textures versus a normal distribution of responses we first computed the mean response to each texture and then computed the non-parametric skew for each neuron across the textures:

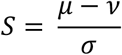

Where μ is the mean, ν the median, and σ the standard deviation.

### Principal components analysis

As previous studies of S1 have quantified the dimensionality of texture representation, repeating these analyses allows for a simple comparison between S1 and S2/PV. Thus, we performed PCA on the mean texture-induced response during the same steady state 300 ms window as described above. PCA was performed on either the shared textures between datasets (**Figure 4**, textures = 38) or on their full respective datasets (**Supplementary Figure 2**). Due to the similar numbers of neurons (SC = 141, S2/PV = 120) we did not control for the number neurons, thus providing a relatively conservative estimate of the dimensionality of S2/PV. We also correlated the eigenvalues from the first 10 dimensions of the PCA analysis with the roughness values from the human psychophysical dataset and squared the result to get the coefficient of determination.

### Population similarity

We also sought to quantify how similar pairs of neurons were to each other. Therefore, we computed the coefficient of determination for all pairs of neurons within each dataset and then took the mean across all pairs to determine the average pairwise similarity.

### Synthetic populations

Given that both the range of firing rates was lower, and the variance was higher in S2/PV, it became obvious that more trials were necessary for 5-fold cross-validated classification to be attempted. Thus, we sought to create a synthetic population for both S2/PV and S1 so that the number of trials could be matched across populations. In S2/PV we implemented a model in which for each neuron we computed the relationship between the mean and variance to the responses across textures. For each neuron we were then able to use the observed mean firing rate for a texture and produce a distribution of firing rates for that texture that matched the mean-variance relationship of that neuron. We found that drawing 10 trials from the distribution was sufficient to overcome the observed issues with 5-fold cross-validation. Comparing the synthetic population against the original data (matching the number of trials) showed that classification performance on the synthetic responses was marginally higher than that of the real data, however the real performance was within the error margin of the synthetic population’s performance.

When attempting to repeat this in SC, however, we found that there was no relationship between the mean firing rate and variance around the mean across textures. Consequently, we instead computed the standardized residual firing rates across all textures:

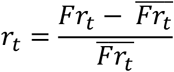

We then fit a gamma distribution to the residuals for each neuron. From there we could again simulate firing rates for each texture by drawing residuals from the fitted distribution and then reversing the above formula.

### Rate classification – nearest neighbor

To determine how well individual neurons could act as classifiers we used cross-validated nearest neighbor classification to quantify both S1 and S2/PV performance. Briefly, we computed the in-fold mean firing rate for each texture for all trials except one (the out of fold trial). We then computed the Euclidean distance between each held-in mean and the held-out firing rate from all textures, selecting the texture with the shortest distance as the predicted texture. We repeated this for all folds (n=5) and computed the proportion of times the shortest Euclidean distance correctly chose the same texture as the held-in trials.

### Rate classification – linear discriminant analysis

To extend the analysis across multiple cells, we implemented 5-fold cross-validated linear discriminant analysis (MATLAB’s *fitdiscr* function) for classification. For each number of neurons, we randomly selected N neurons (without replacement) from the population and performed the classification. This was repeated 100 times for each N to build an estimate of the distribution of performance for each value of N across the population.

### Sliding window rate classification

In addition to performing LDA classification on texture identity during the steady-state period of the initial response, we also tested the extent to which the neural response was informative about both the trial type and the animal’s decision. To do this, we repeated the previous LDA method but with a 250 ms window that was incremented by 50 ms over the duration of the trial. This allowed us to determine when the respective information became present, if at all. In both cases we subsampled 25 trials from each condition (same vs different trial or animal chose same vs animal chose different) and attempted to classify the appropriate outcome, again using 5-fold cross validation.

For the trial-type classification implementation we observed that we were able to classify trial type at the onset of the first stimulus. This led us to identify several textures that were inhomogeneous across the scanning direction allowing for the classifier to determine trial type depending on whether the first or second half of the texture was presented during the first interval. Consequently, for that analysis we removed 13 textures that contributed to this effect.

### Roughness decoding

Given the relatively poor performance of S2/PV neurons as classifiers, we sought to quantify how much textural information was contained by attempting to decode roughness – the dominant texture property - from the responses. Thus, we performed cross-validated regressions using the mean response to each texture and the psychophysical ratings. First, we took a random sample of 15 neurons from the population, then we fit a regression model on the firing rates of those neurons against the roughness ratings for all textures except a held-out texture, finally we predicted the roughness of the held-out texture using the same model. Once this was repeated for all textures, we then correlated the predicted roughness values with the reference ratings. This was repeated 1000 times, where on each permutation a different sample of 15 neurons was chosen.

### Timing classification

The timing analysis performed here has been described in more detail previously (Long et al., 2022). Briefly, for individual neurons smoothing filters of different widths were applied to the steady state response of each trial. For a given smoothing window, the mean maximum cross-correlation was computed for each pair of textures across all trials. Then, nearest neighbor classification was performed where the correlation value was treated as the distance and the class with the greatest correlation was selected. This was repeated for all smoothing windows and the window with the greatest performance was chosen for comparison with the rate-based nearest neighbor classifier.

## Supplementary Figures

**Supplementary Figure 1.**
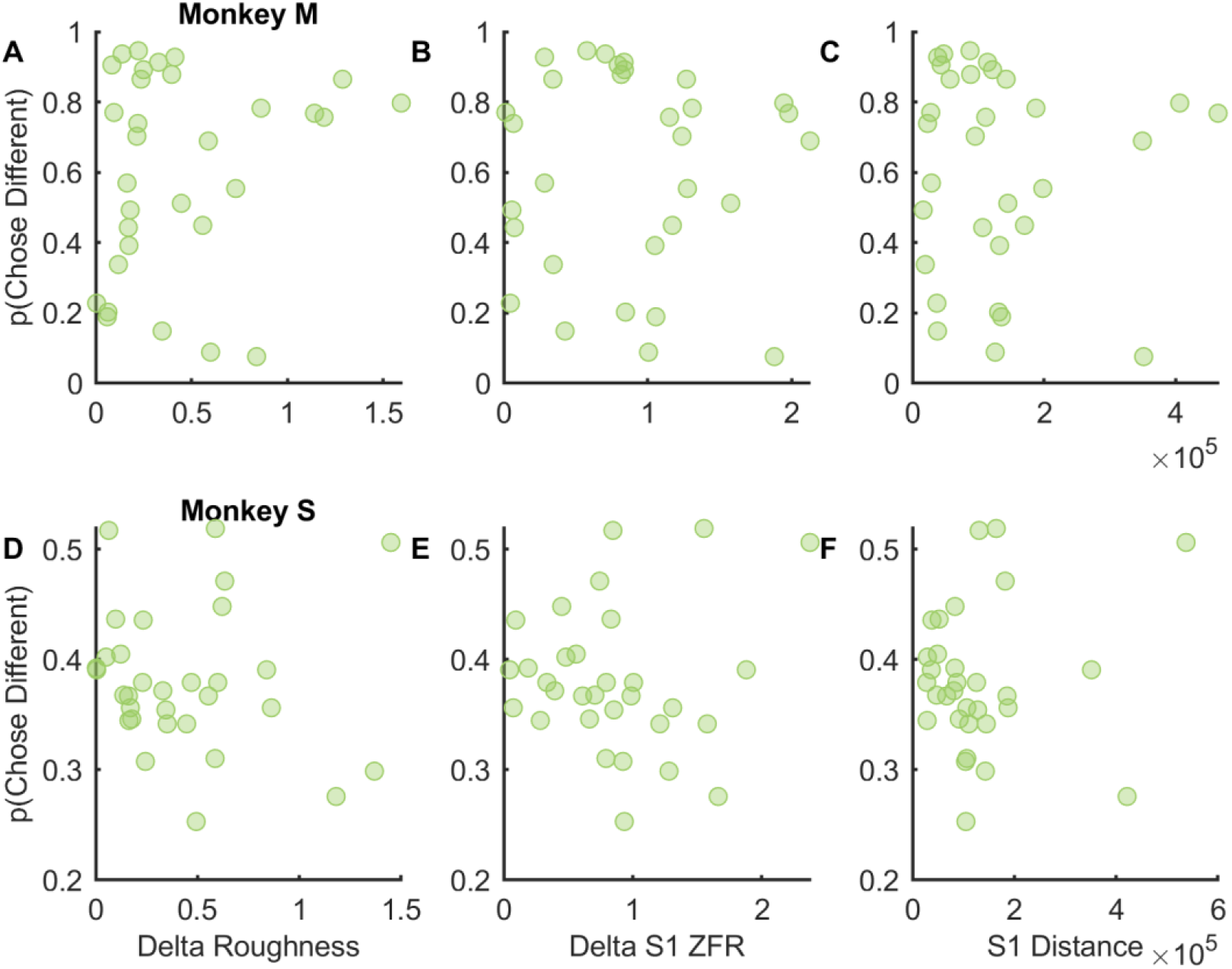
Expected predictors of the animal’s decision are poor. A, D| The difference in roughness for a given pair of textures does not correlate with the animal’s likelihood of choosing different. B, E| The population-wide difference in firing rate (Z-normalized firing rate across neurons) also does not predict decision rates. C, F| The Euclidean distance between pairs of textures in S1 space is not a predictor of decision rates. R^2^ < 0.04 for all comparisons.

**Supplementary Figure 2.**
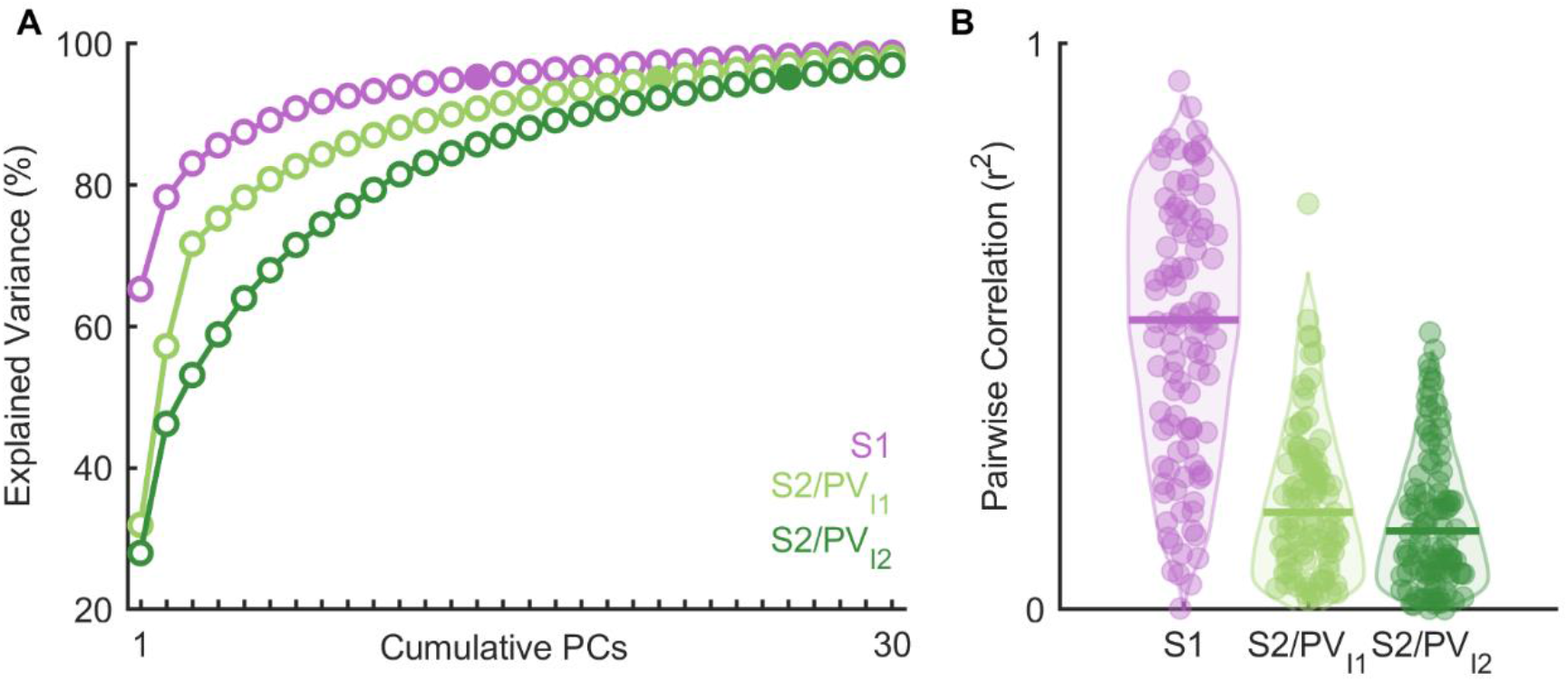
|Dimensionality of responses on full datasets. A| Cumulative explained variance with respect to principal component for SC, and S2/PV (both interval 1 and interval 2). The second interval is higher dimensional than the first interval. B| Pairwise correlation values between all pairs of neurons for each population similarly shows that S1 neurons tend to be much more similar to one another than those in S2/PV during either interval.

**Supplementary Figure 3.**
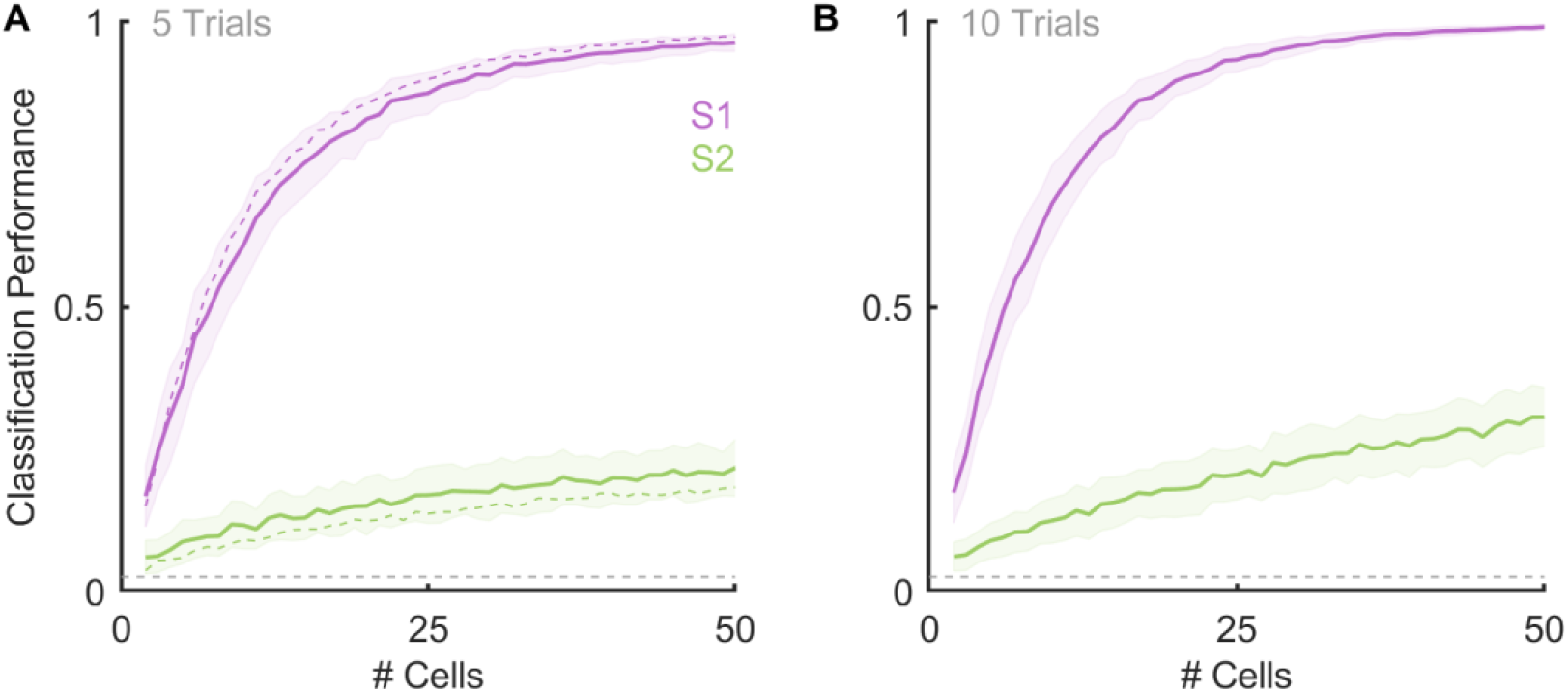
Linear discriminant analysis using the responses of a synthetic population. A| Classification performance using the responses of the synthetic population (solid lines) or the original data (dashed lines) using 5 trials for both (the number present in the original dataset; 5-fold cross-validation). Shaded areas represent standard deviation. B| Classification performance using the responses of the synthetic population when the number of trials is increased to 10 (5-fold cross-validation).

**Supplementary Figure 4.**
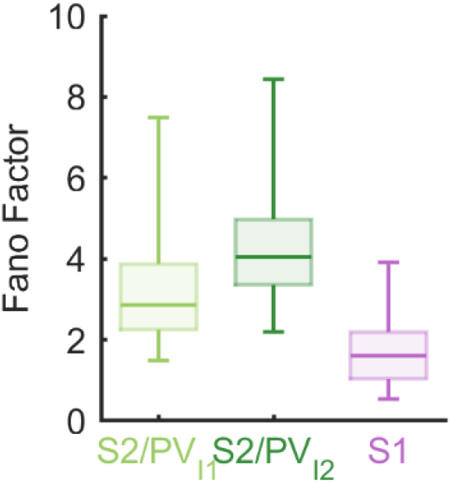
The Fano factor of S2/PV responses is much higher than its S1 counterpart.

## Notes

### Competing Interest Statement

The authors have declared no competing interest.

### Summary of Updates

Edits throughout. Rewrite the discussion.

